# Reduced SK channel control of mesolimbic dopamine neuron firing drives reward seeking adaptations in chronic pain

**DOI:** 10.1101/2025.10.06.680596

**Authors:** Jeremy M. Thompson, Robert D. Graham, Jeff P. Goff, Yu-Hsuan Chang, Vani Kalyanaraman, Chang Xu, Mason Barrett, Hanyun Wang, Travis Hage, Alex Legaria, Mengtong Liu, Rebecca George, Bryan A. Copits, Alexxai Kravitz, Meaghan C. Creed

## Abstract

Patients with chronic neuropathic pain typically experience affective symptoms that drive reduced quality of life and negatively impact pain management. Mesolimbic dopamine is necessary for reward valuation and learning, and the existence of a hypodopaminergic state has been proposed to underlie these affective symptoms of chronic pain. However, direct functional evidence for this hypodopaminergic state is lacking, and the mechanisms underlying its emergence over the acute to chronic pain transition are unknown. Here, we find a selective deficit in the ability of mesolimbic dopamine neurons to sustain burst firing, which is apparent uniquely at chronic timepoints following neuropathic injury. As a result, animals are unable to sustain effortful pursuit of rewards under conditions of high effort or time costs. Convergent biophysical modeling and experimental electrophysiology establish that in a spared nerve injury (SNI) model of chronic neuropathic pain, calcium-activated, small-conductance potassium (SK) channel function is impaired, resulting in lower peak firing and earlier entry into depolarization block of mesolimbic dopamine neurons. Critically, dopamine dependent reward learning, formation of cue-reward associations and locomotor activity remain intact, arguing against the interpretation of a generalized hypodopaminergic state. These results elucidate a circuit-level basis for selective motivational deficits emerging in chronic neuropathic pain.

## Introduction

Chronic neuropathic pain afflicts millions of people worldwide, yet remains notoriously difficult to treat^1^. A major obstacle is the frequent emergence of comorbid depressive symptoms that amplify the perception of pain^2^, impair quality of life^3-6^ and reduce patient compliance with pain management therapies^2-6^. These negative affective symptoms are typically refractory to conventional antidepressant therapies^7-10^, compounding clinical burden and underscoring a key gap in our understanding of how pain alters reward processing. Elucidating the cellular and neurocircuit mechanisms driving the emergence of depressive symptoms is needed to develop new strategies to manage chronic pain.

Negative affect in chronic pain has been proposed to be linked to hypodopaminergia^11-14^. Patients with chronic neuropathic pain exhibit altered activity and connectivity of the ventral tegmental area^15-17^, reduced ventral striatal dopamine (DA) transporter binding^16-18^ and dopamine Drd2 receptor binding^19, 20^, consistent with DA dysfunction. However, whether these changes are a consequence of pain, reflect compensatory adaptations to changes in DA signaling or are antecedent to pain and increase CNP susceptibility, cannot be resolved with cross-sectional patient studies. Mechanistic studies of pain-related hypodopaminergia in rodents have been paradoxical, with blunted basal DA levels^21-23^, and increased phasic dopamine signaling in response to rewarding^23, 24^ or aversive^21^ stimuli often, but not always^25^ reported. Critically, these studies have overwhelmingly examined *acute* pain states (within 10 days of nerve injury), failing to capture chronic adaptations^23, 26-28^. Of studies that examined VTA activity several weeks following injury, firing of DA neurons was either *increased*, or heterogeneous effects were observed, depending on anatomical and electrophysiological subclass^29-32^. Despite the extensive interest, the time course, cellular mechanisms and causal relationship between chronic pain, mesolimbic dopamine adaptations and negative affect remains unresolved.

Here, we measured behavior on multiple tasks that are known to be dependent on mesolimbic DA function, and identified selective impairments in reward valuation, while reward learning and behavioral flexibility remained intact in chronic pain states. Excitability and basal firing of mesolimbic DA neurons was also unchanged in chronic pain, however neurons were unable to sustain consistent action potential firing due to earlier entry into depolarization block. Critically, both the behavioral and electrophysiological adaptations were only evident at chronic timepoints post nerve injury. Convergent computational modeling and electrophysiology identified reduced small conductance, calcium-activated potassium (SK) channel function as a key driver of this electrophysiological phenotype; and restoring SK function rescued pain-induced adaptations in both mesolimbic DA firing and reward valuation following chronic spared nerve injury (SNI). Together, our results establish a causal relationship between altered biophysical properties of mesolimbic DA neurons and changes in reward valuation in chronic pain, and point to SK channels as a potential entry point for targeting affective symptoms of chronic neuropathic pain.

## Results

### Reward-guided learning is preserved in a model of chronic neuropathic pain

We used reward-guided decision-making tasks to quantify changes in reward sensitivity, operationally defined as the ability to update behavior in response to rewards^33^. To avoid experimental confounds of food restriction or physical requirements which interact with pain states^34^, we used closed economy tasks to measure reward-learning over multiple days in the homecage using FED3.0 (**Fig 1A**)^35^. SNI mice exhibited consistent allodynia for up to 8 weeks (**Fig 1B**). There was no difference between SHAM and SNI mice in the ability to learn the FR1 contingency to nosepoke to earn a food pellet (**Fig 1C**), and groups earned equivalent pellets from the FED3.0 across all tasks (**Fig 1D,E**). Surprisingly, there were no differences between SHAM and SNI mice in their accuracy on either deterministic (100:0) or probabilistic (80:20) versions of a reversal learning task, or on their ability to use prior reward history to guide decision making (**Fig 1F-I**).The lack of differences could be because in both deterministic and probabilistic reversal tasks, the structure is relatively constrained, without adequate volatility to reveal differences in decision making strategy. So, we next tested mice in a dynamic foraging task, where the overall reward rate in the environment could be high, medium or low, with a 30% difference in probabilities between the two nose poke options (**Fig 1J**). Again, we found no difference in pellets earned from the environment, or in integration of reward history or accuracy between SHAM and SNI mice (**Fig 1K-N**). However, when the reward rate in the environment was high, SHAM mice were faster to initiate a new trial, while SNI mice performed equivalently across blocks, indicating a lack of sensitivity to reward availability in SNI mice (**Fig 1O**). This lack of reward sensitivity was not due to general slowness in the task, since pellet retrieval time was slightly but significantly faster in SNI mice (**Fig 1P**).

**Figure 1.**
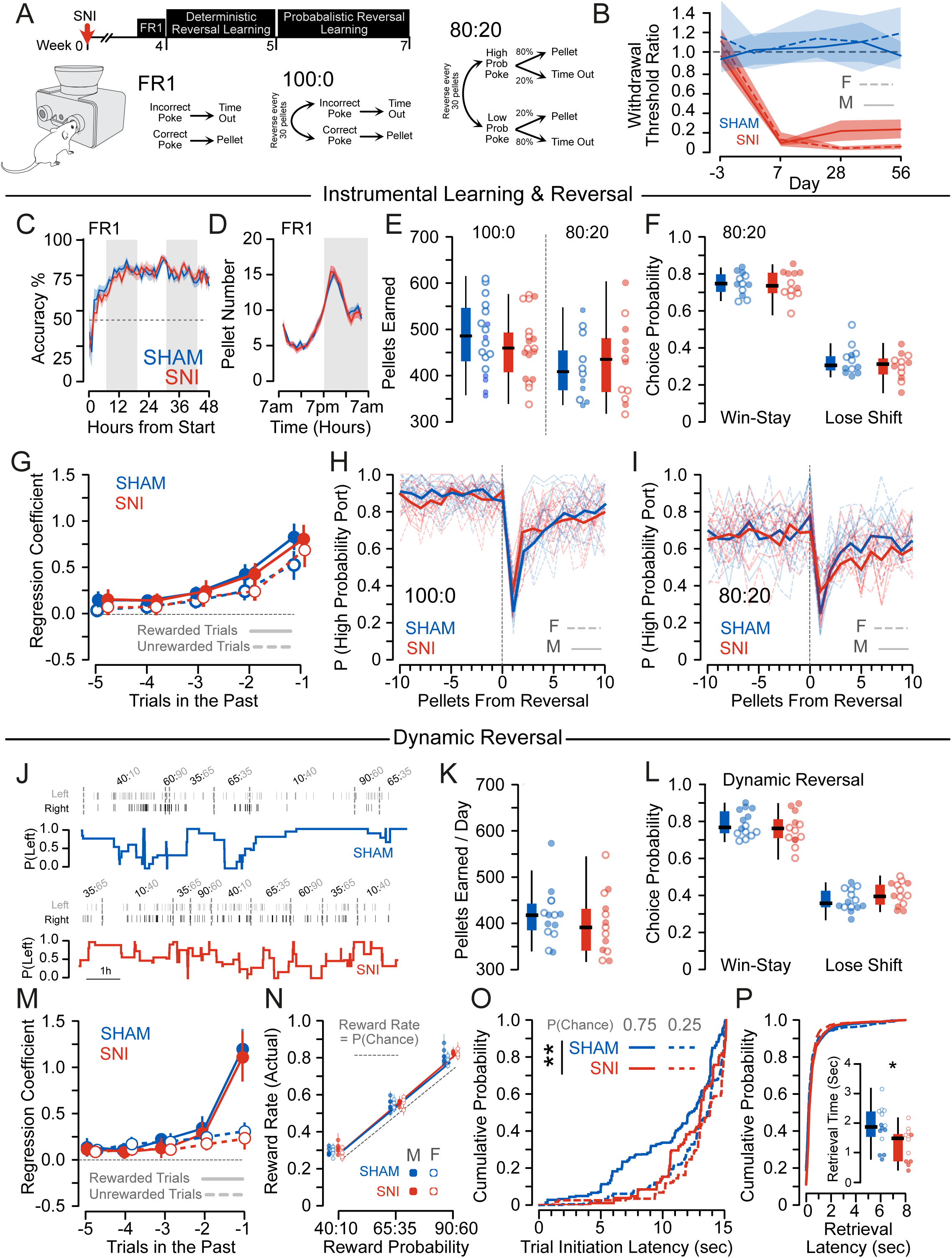
Reward-guided decision making is preserved following chronic nerve injury. (A) Experimental timeline and schematic of task design. (B) Mechanical withdrawal thresholds as measured with the von Frey assay; mechanical hypersensitivity persisted at 8 weeks post-SNI (Baseline: 1.08 ± 0.05, 8 weeks post: 0.19 ± 0.03, F_Timepoint_ = 178.962, p < 0.001, SHAM n = 10M/7F, SNI n = 18M/18F). (C) There was no difference in FR1 task acquisition between SHAM and SNI mice (F_Group_ = 1.175, p = 0.280). (D) There was no difference in number or in pattern of pellets earned on the FR1 task between SHAM and SNI mice (F_Group_ = 0.678, p = 0.412). (E) There was no difference in pellets earned in the first 72h of deterministic (100:0) or probabilistic (80:20) reversal learning tasks. (Deterministic: SHAM = 532.16 ± 22.38, SNI = 506.06 ± 21.27, F_Pellet_ = 0.712, p = 0.404, n_SHAM_ = 6M/12F, n_SNI_ = 6M/12F; Probabilistic: SHAM = 448.25 ± 24.419, SNI = 464.42 ± 31.435, F_Pellet_ = 0.165, p = 0.689, n_SHAM_ = 7M/7F, n_SNI_ = 6M/7F). Males: closed circles, females: open circles. (F) There was no difference in Win-stay (SHAM = 74.38 ± 1.75, SNI = 74.36 ± 2.31, t = 0.01, p = 0.996) or in Lose-shift (SHAM = 32.96 ± 2.27, SNI = 30.10 ± 2.28, t = 0.886, p = 0.385) strategy on the probabilistic reversal task between SHAM and SNI mice. Males: closed circles, females: open circles. (G) Logistic regression revealed significant decay of weighting of rewarded (F_Trial_=144.46, p<0.001) and unreward (F_Trial_ = 102.59, p < 0.001) choices over time, but no difference in integration of prior reward history between SHAM and SNI mice on the probabilistic reversal task (rewarded: F_Group_ = 0.020, p = 0.886, F_Interaction_= 0.272, p = 0.992, unrewarded: F_Group_ = 0.299, p = 0.585, F_Interaction_= 4.285, p = 0.369). (H) There were no differences between SHAM and SNI mice on accuracy in the deterministic reversal task (F_group_ = 0.498, p = 0.485). (I) There were no differences between SHAM and SNI mice on the probabilistic reversal task (F_group_ = 0.366, p = 0.718). (J) Representative performance on the dynamic foraging task. (K) There was no difference in pellets earned in the first 72h of the dynamic foraging task (SHAM = 390.36 ± 15.1, SNI = 405.92 ± 19.22, t = 0.65, p = 0.525, n_SHAM_ = 7M/7F, n_SNI_ = 6M/7F). Males: closed circles, females: open circles. (L) There was no difference in Win-stay (SHAM = 79.65 ± 4.22, SNI = 76.98 ± 2.91, t = 0.85, p = 0.401) or in Lose-shift (SHAM = 38.15 ± 0.42, SNI = 41.16 ± 3.64, t = 1.26, p = 0.218) strategy on the dynamic foraging task between SHAM and SNI mice. Males: closed circles, females: open circles. (M) Logistic regression revealed significant decay of weighting of rewarded (F_Trial_=178.88, p<0.001) and unreward (F_Trial_= 50.725, p < 0.001) choices over time, but no difference in integration of prior reward history between SHAM and SNI mice on the probabilistic reversal task (rewarded: F_Group_ = 0.979, p = 0.886, F_Interaction_= 0.272, p = 0.322, unrewarded: F_Group_ = 1.657, p = 0.198, F_Interaction_= 3.000, p = 0.558). Males: closed circles, females: open circles. (N) Mice performed significantly better than chance across all environmental reward rates (SHAM; t = 11.213, p < 0.001, SNI: t = 12.473, p < 0.001), without significant differences between groups (t = 1.314, p = 0.189). (O) SHAM mice exhibited a lower trial initiation latency in High Reward (90:60) blocks relative to Low Reward (40:10) blocks (p = 0.017), this difference was absent in SNI mice (p = 0.157; SHAM vs. SNI: t=2.594, p = 0.010) (P) SNI mice exhibit faster pellet retrieval times relative to SHAM mice (SHAM = 1.85 ± 0.18 sec, SNI = 1.27 ± 0.16 sec, t = 2.35, p = 0.027). Males: closed circles, females: open circles.

### Mice expend less effort in pursuit of rewards in chronic neuropathic pain

To test whether the observed deficit in sensitivity reflects changes in reward motivation, we tested the willingness of mice to work for rewards on a progressive ratio (PR) task, where pellet cost incremented with each successive pellet, with a reset after 30 minutes of inactivity (**Fig 2A-B**). Body weight and pellets earned from the task were equivalent between groups (**Fig 2C, D, G**). However, mice exhibited reduced willingness to work for rewards following chronic nerve injury; SNI mice exhibited significantly shorter runs before reaching break points (**Fig 2E**), and expended significantly fewer nosepokes to earn equivalent pellets from the task (**Fig 2F**). Critically, these differences were not apparent acutely (within 1 week) following SNI (**Fig. 2G-I**), suggesting this phenotype is unique to the chronic pain state. Corroborating this interpretation, there were no changes in pokes per pellet or task performance following acute paw incisional pain (**Fig S1A-G**). Moreover, once established in the chronic SNI mice, transient alleviation of hyperalgesia with metformin (**Fig S1H**) did not change activity or strategy on the PR task (**Fig S1I-O**), indicating that the changes in reward valuation are dissociable from the sensory symptoms following SNI. We also confirmed that deficits in PR performance were not due to reduced task engagement due to motor effects, since homecage locomotor activity was not different between groups (**Fig 2J**).

**Figure 2.**
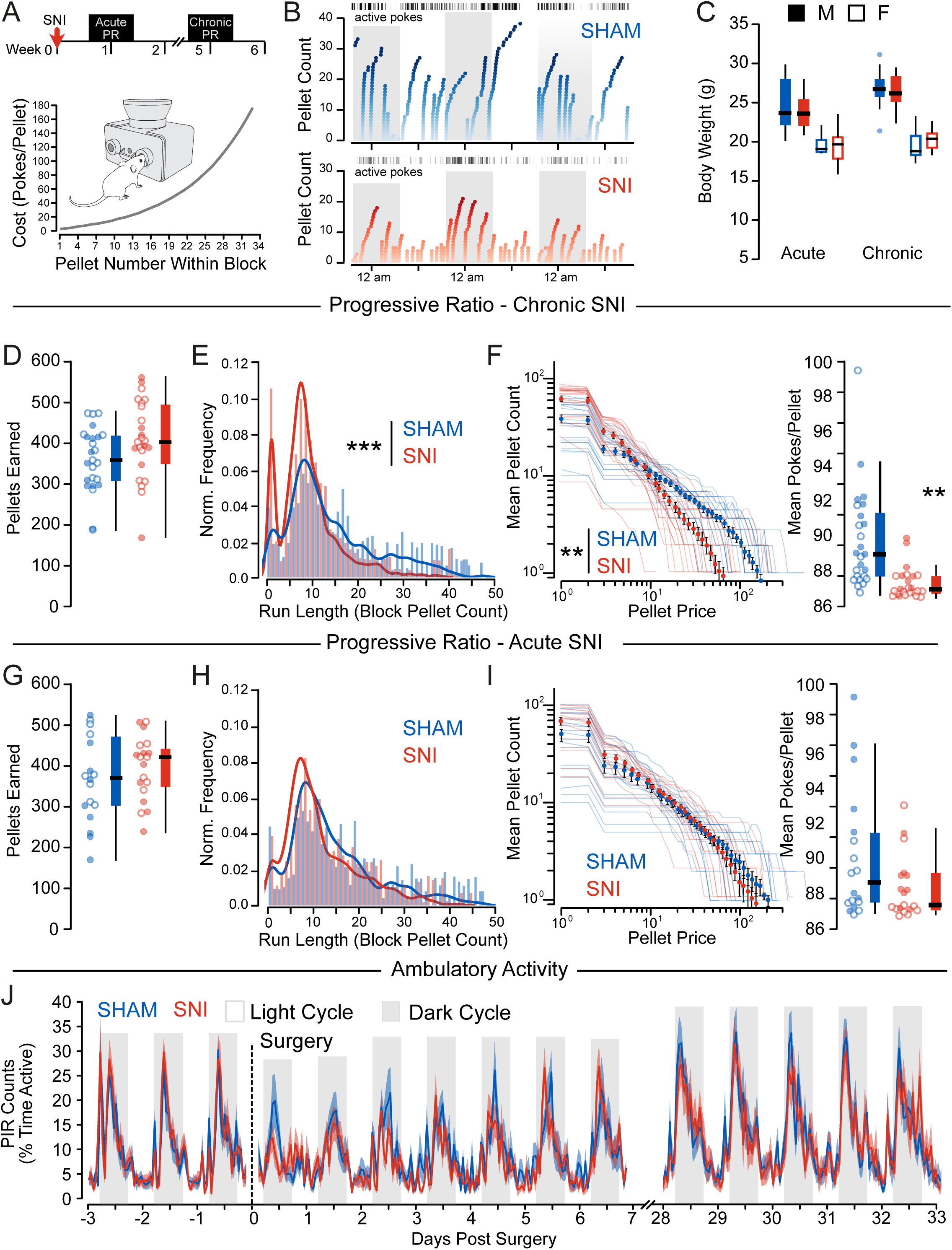
Mice adopt an efficient reward-seeking strategy following chronic spared nerve injury. (A) Experimental timeline and schematic of resetting progressive ratio (PR) task showing increased pokes required to earn successive pellets within a block; effort requirement reset to 1 poke after 30 minutes of inactivity. (B) Representative examples of SHAM (top) and SNI (bottom) animals performing the rPR task over three days, dark cycle is shaded. (C) There were no differences in body weight between the SHAM and SNI groups at either the acute (F_Group_ = 1.95, p=0.172) or chronic (F_Group_ = 0.26, p=0.615) timepoint (Acute: n_SHAM_ = 9M/9F, n_SNI_ = 9M/9F, Chronic: n_SHAM_ = 11M/14F, n_SNI_ = 9M/15F) (D) At chronic timepoints following SNI, there was no difference in pellets earned (F_Group_ = 1.92, p = 0.17). Males: closed circles, females: open circles. (E) SNI mice exhibited lower progressive ratio run lengths (KS: 0.241, p<0.001). (F) SNI mice expended fewer pokes per pellet earned (F_Group_ = 5.62, p = 0.005). (G) There was no difference in pellets earned acutely post-SNI (F_Group_ = 1.41, p = 0.24). Males: closed circles, females: open circles. (H) There was no significant difference in the run lengths (KS: 0.045, p = 0.2422) acutely post-SNI between SHAM and SNI mice. (I) There was no significant difference in pokes per pellet (F_Group_ = 1.92, p = 0.17) acutely post-SNI between SHAM and SNI mice. (J) Patterns of homecage locomotor activity were not significantly different between SHAM and SNI groups acutely (F_Group_ = 0.42, p = 0.527, F_Time_ = 7.61, p > 0.001, F_GroupxTime_ = 1.16, p = 0.082) or 28 days post-injury (F_Group_ = 0.01, p = 0.938, F_Time_ = 9.61, p > 0.001, F_GroupxTime_ = 0.70, p = 0.999).

### Mesolimbic dopamine neurons fail to sustain firing following chronic SNI

The selective impairment SNI mice exhibited in effort-based decision making with preservation of reward-guided learning is reminiscent of behavioral changes with dopamine antagonism (i.e. haloperidol; **Fig S2**). In rodents, the ventral striatum is divided into two distinct subregions: the nucleus accumbens core (NAcC) and shell (NAcSh), which are innervated by separate populations of DA neurons in the VTA^36, 37^. DA release in the NAcC (*but not NAcSh*) reflects the anticipated value of upcoming rewards^38-42^ and is critical for maintaining effortful pursuit of rewards^43-45^. Therefore, we used retrograde labeling to identify and record from NAcC-projecting DA neurons in the VTA (mesolimbic DA neurons; **Fig 3A**). Consistent with prior work, NAc core-projecting DA neurons were located within the paranigral nucleus of the VTA, and expressed the calcium-binding marker calbindin^37, 46^ in addition to tyrosine hydroxylase (**Fig 3B-C**). Relative to naive mice or SHAM controls, SNI mice exhibited significantly reduced evoked action potential firing (**Fig. 3D-E**) with an increase in interspike intervals at peak firing frequency (**Fig. 3F**) and without changes in rheobase (**Fig. 3G**). This altered firing was characterized by earlier entry into depolarization block in mesolimbic DA neurons from SNI mice (**Fig. 3H**), resulting in reduced peak firing rate (**Fig. 3I**). Interestingly, despite reduced firing, changes in intrinsic membrane properties in SNI mice were consistent with increased excitability (**Fig. S3**); although resting potential (**Fig S3A**) and spontaneous firing rate (**Fig. S3B**) were unchanged, SNI neurons exhibited significantly increased input resistance relative to naive controls (**Fig. S3C**). Critically, these changes in action potential firing were not present at an acute (1 week) timepoint following SNI (**Fig. 3J-O**, **Fig. S3E-H**), suggesting that these changes emerge with the transition from acute to chronic pain.

**Figure 3.**
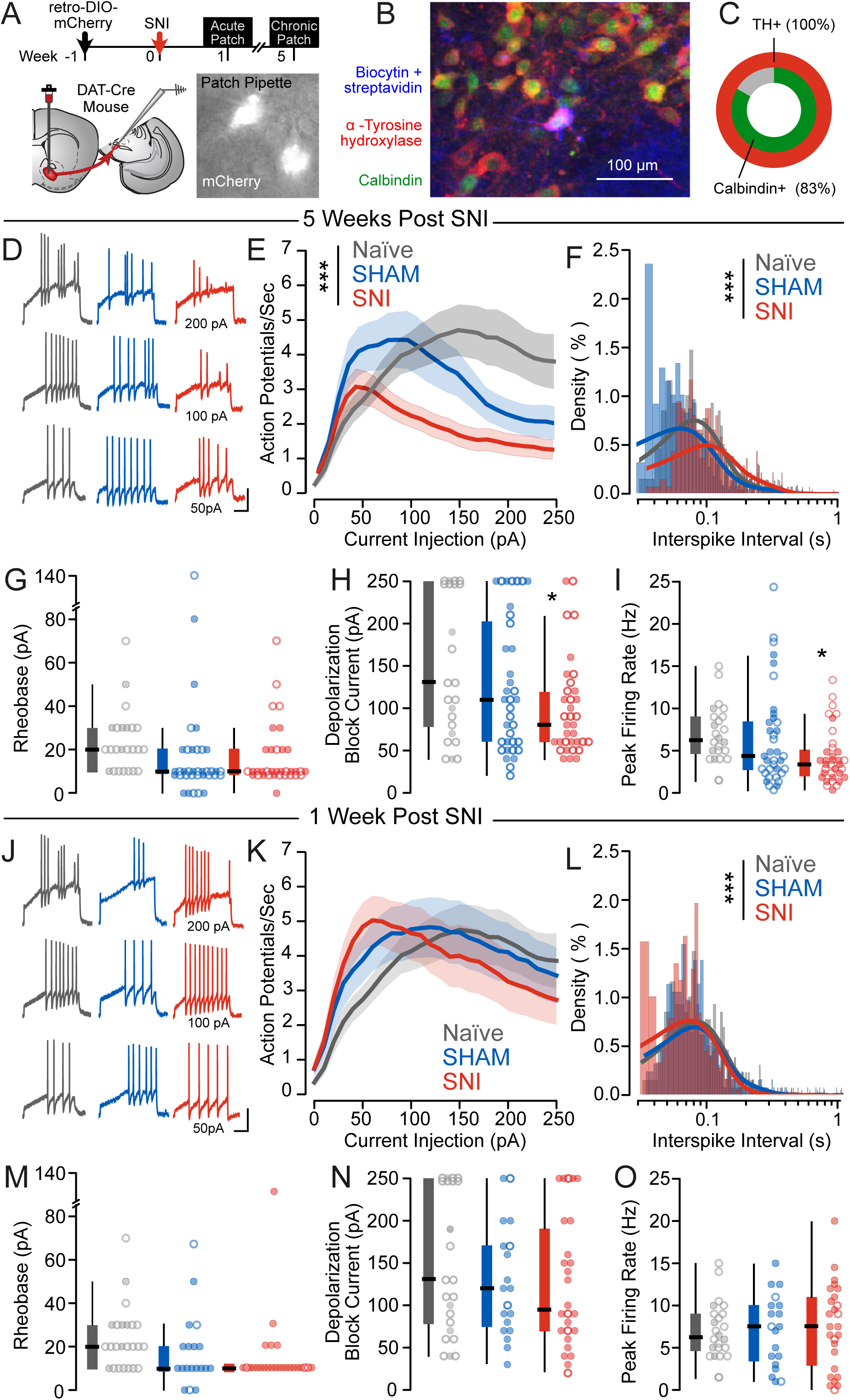
Depolarization block prevents sustained firing of mesolimbic dopamine neurons after chronic SNI. (A) Experimental protocol for patch clamp electrophysiology experiments. (B) Representative image of post-hoc immunohistochemistry of recorded cells. (C) All recorded neurons characterized by post-hoc immunohistochemistry were positive for tyrosine hydroxylase (18/18 cells) and 83% (15/18 cells) were positive for calbindin. (D) Representative traces (scale bar: 20 mV, 500 ms) of action potential firing following stepwise current injections are shown for SHAM and SNI neurons. (E) SNI neurons (n_SNI_ = 19M/16F, N_SNI_ = 9M/7F, n_SHAM_ = 16M/20F, N = 9M/11F) had reduced action potential firing (F_Group_ = 31.07, p_Group_ < 0.0001, F_Sex_ = 11.87, p_Sex_ = 0.0006) chronically after SNI. (F) Interspike interval distribution at the peak firing frequency was increased in SNI neurons (KS_NAIVE-SHAM_ = 0.22, p_NAIVE-SHAM_ < 0.0001, KS_NAIVE-SNI_ = 0.20, p_NAIVE-SNI_ < 0.0001, KS_SHAM-SNI_ = 0.36, p_SHAM-SNI_ < 0.0001). (G) Rheobase was unchanged (F_Group_ = 0.81, p_Group_ = 0.45, F_Sex_ = 0.12, p_Sex_ = 0.73) following SNI at a chronic timepoint. Males: closed circles, females: open circles. (H) SNI neurons entered into depolarization block at lower currents (F_Group_ = 3.24, p_Group_ = 0.04, F_Sex_ = 0.05, p_Sex_ =0.82) than SHAM or NAIVE mice at a chronic timepoint after surgery. Males: closed circles, females: open circles. (I) There was a trend for reduced peak firing rate in response to depolarizing current steps in SNI neurons (F_Group_ = 2.57, p_Group_ = 0.08, F_Sex_ = 0.82, p_Sex_ = 0.37). Males: closed circles, females: open circles. (J) Representative traces (scale bar: 20 mV, 500 ms) of action potential firing following stepwise current injections are shown for SHAM and SNI neurons. (K) There was no difference in action potential firing in response to 2 s depolarizing current steps acutely following SNI (F_Group_ = 1.25, p_Group_ = 0.29, F_Sex_ = 3.65, p_Sex_ = 0.06, n_SNI_ = 22M/4F, N_SNI_ = 10M/2F, n_SHAM_ = 16M/3F, N_SHAM_ = 6M/1F, n_NAIVE_ = 6M/18F, N_NAIVE_ = 2M/5F). (L) There was a small decrease in interspike intervals at the peak firing frequency at an acute 1 week timepoint following SNI (KS_NAIVE-SHAM_ = 0.10, p_NAIVE-SHAM_ = 0.14, KS_NAIVE-SNI_ = 0.23, p_NAIVE-SNI_ < 0.0001, KS_SHAM-SNI_ = 0.18, p_SHAM-SNI_ = 0.0001). (M) There was no difference in rheobase at an acute 1 week timepoint following SNI (F_Group_ = 0.58, p_Group_ = 0.57, F_Sex_ < 0.01, p_Sex_ = 0.98). Males: closed circles, females: open circles. (N) There was no difference in current at entry into depolarization block acutely following SNI (F_Group_ = 0.24, p_Group_ = 0.79, F_Sex_ = 0.26, p_Sex_ = 0.61). Males: closed circles, females: open circles. (O) There was no difference in peak firing rate at an acute 1 week timepoint following SNI (F_Group_ = 0.07, p_Group_ = 0.94, F_Sex_ = 0.09, p_Sex_ = 0.76). Males: closed circles, females: open circles.

### Dopamine neuron hypoactivity following chronic SNI is related to failure of sodium channel recovery from inactivation due to SK channel dysfunction

To investigate the mechanisms underlying the pain-related firing adaptations, we developed multicompartment cable models of mesolimbic DA neurons in SNI and SHAM conditions. We used a Markov Chain Monte Carlo (MCMC) method constrained by the experimentally measured rheobases, input-output curves, medium afterhyperpolarization potentials, and input resistances (**Fig. 4A-D**). The MCMC method produced 50 combinations of maximal ion channel conductances describing SNI neurons and 64 combinations describing SHAM neurons. The SNI and SHAM model populations predicted decreased peak voltage-gated sodium channel (NaV1.2) conductance (**Fig. 4E**), increased transient, rapidly activating and inactivating voltage-gated potassium (A-type) conductance (**Fig. S4A**), and decreased SK channel conductance (**Fig. 4I**) in mesolimbic dopamine neurons following SNI, which we then validated experimentally. To evaluate NaV function, we examined action potentials evoked by a depolarizing current ramp. The first action potential formed did not exhibit differences in threshold or phase plot (**Fig. 4F-G**), however the distribution of subsequent action potential heights was decreased in SNI neurons relative to SHAM controls (**Fig. 4H**). This suggests that SNI neurons initially fire normally, but cannot sustain repeated firing due to inability to recover from inactivation. SK channels are critical regulators of dopamine neuron firing and play an important role in sodium channel recovery from inactivation^48-50^, consistent with our model prediction of reduces SK conductance in SNI (**Fig. 4I**). We tested this prediction by measuring action potential half-width and the magnitude of the mAHP formed after depolarizing current steps of increasing magnitude which revealed widening of the action potential and reduced mAHP amplitude in SNI neurons (**Fig. 4J-K**). We confirmed decreased SK currents by measuring the outward current formed following depolarizing voltage steps before and after selective SK channel blockade (apamin, 100 nM, **Fig. 4L**).

**Figure 4.**
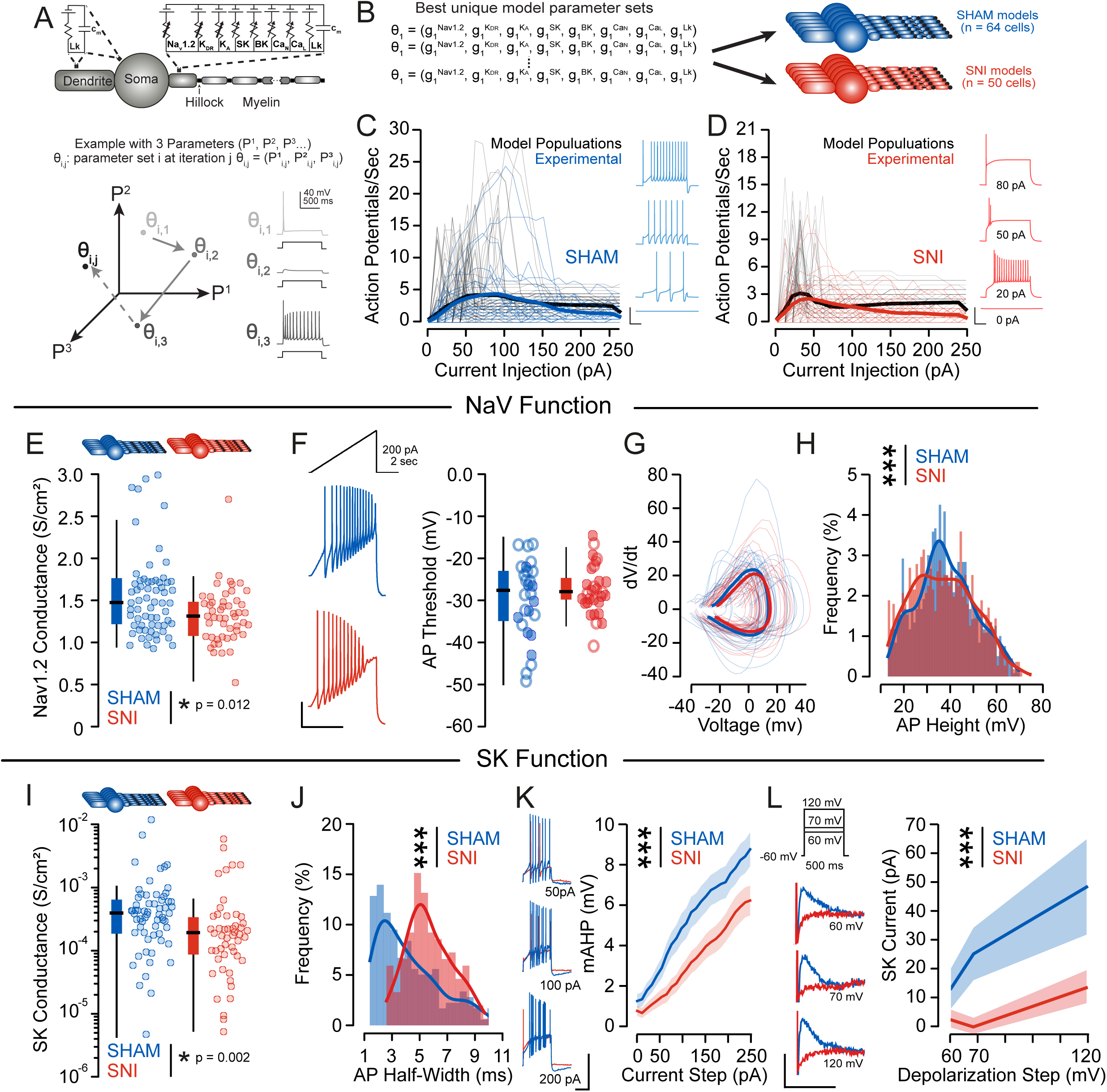
Changes in SK channel function drive altered firing properties of mesolimbic dopamine neurons following chronic SNI. (A) Schematic multicompartment model containing explicit representations of the voltage- and calcium-gated ion channels found in mesolimbic dopamine neurons (top). MCMC method to parametrize populations of biophysically plausible mesolimbic dopamine neurons. (B) The MCMC method produced 64 combinations of maximal ion channel conductances that described SHAM neuron models, and 50 combinations of maximal ion channel conductances that described SNI neuron models. (C) Comparison of SHAM model and experimental neurons I/O curves and traces of model neuron firing (insets). Thin lines represent individual neurons’ I/O curves, solid lines are population means. Black lines represent model neurons, blue lines represent experimentally recorded neurons. (D) Comparison of SNI model and experimental neurons I/O curves and traces of model neuron firing (insets). Thin lines represent individual neurons’ I/O curves, solid lines are population means. Black lines represent model neurons, red lines represent experimentally recorded neurons. (E) SNI neuron models had significantly lower Nav1.2 conductance (1.335 ± 0.046 S/cm^2^) than SHAM neuron models (1.558 ± 0.060 S/cm^2^) (N_SNI_ = 50, N_SHAM_ = 64, U = 1160, p = 0.012) (F) Representative responses to experimental ramp current injections (inset). Action potential threshold was not different between SHAM: -30.5 mV +/- 1.76, and SNI: -29.6 +/- 1.00 mV, F = 1.63, p = 0.21. Males: closed circles, females: open circles. (G) The phase plot of the first action potential formed from a ramp depolarization protocol was unchanged by SNI (n_SNI_ = 13M/11F, N_SNI_ = 8M/5F, n_SHAM_ = 7M/16F, N_SHAM_ = 6M/9F, Max |t| = 12.04, p = 0.45) (H) The action potential height measured at peak firing rate was significantly reduced following SNI (KS = 0.094, p < 0.001) (I) SNI neuron models had significantly lower SK conductance (0.566 ± 0.166 mS/cm^2^) than SHAM (0.840 ± 0.230 mS/cm^2^) neuron models (N_SNI_ = 50, N_SHAM_ = 64, U = 1062, p = 0.002). (J) The action potential half-width was significantly increased in SNI neurons relative to SHAM (KS = 0.418, p < 0.001). (K) The mAHP was decreased following SNI (n_SNI_ = 19M/16F, N_SNI_ = 9M/7F, n_SHAM_ = 16M/20F, N_SHAM_ = 9M/11F, F_Group_ = 127.48, p < 0.001). Representative traces are shown (scale bars: 50 mV, 1 s) (L) SK channel currents were reduced following SNI (n_SNI_ = 5M/6F, N_SNI_ = 3M/3F, n_SHAM_ = 11M/4F, N_SHAM_ = 4M/2F, F_Group_ = 8.29, p = 0.005). Representative traces are shown (scale bars: 100 pA, 10 ms)

Finally, we also evaluated the model prediction of increased A-type potassium conductance. Because the A-type potassium current activates near the action potential threshold and delays the onset of firing^47^, we first assessed A-type potassium current activity by quantifying the time to the first action potential formed by stepwise current injections. We found that latency of action potential firing was significantly increased in SNI neurons (**Fig. S4B**), however when directly measuring the A-type potassium current in voltage clamp before and after application of a selective blocker (AmmTx3, 10 nM), we did not observe any group differences (**Fig. S4C**). This suggests that increased A-type potassium current is not a primary driver of chronic pain-related hypoactivity.

Together, these results indicate that chronic pain-related mesolimbic DA neuron firing adaptations are driven by failure of sodium channel recovery from inactivation arising from SK channel dysfunction.

### Over-expression of SK channels rescues cellular and behavioral adaptations in the chronic pain state

To investigate causality of SK channel dysfunction on dopamine neuron hypoactivity, we developed a retrograde floxed SK3 subunit-expressing AAV (**Fig. 5A**) to over-express the predominant SK channel subunit^51^ in NAc-projecting VTA neurons to rescue SK channel function. Following injection of this virus into the NAc core, we were able to detect expression of the FLAG tag associated with the virally expressed SK3 subunit (**Fig. 5B**) and found a significant increase in SK channel currents (**Fig. 5C**), confirming expression and function of the virus. This viral-mediated SK3 over expression significantly increased action potential firing (**Fig. 5D**) and normalized current at entry into depolarization block (**Fig. 5E**). Crucially, we saw an increase in action potential height, decrease in action potential half-width, and decrease in interspike intervals (**Fig. 5G-H, Fig. S5C**), indicating that normalization of SK channel function was sufficient to allow sodium channel recovery from inactivation, thereby increasing peak action potential firing (**Fig S5. A-B**). We then investigated behavioral effects of SK3 over-expression on the rPR task (**Fig. 5I-J**). Body weight (**Fig. 5K**) was unchanged four weeks following injection of an SK3-expressing AAV. SK3 AAV-injected mice had a significant increase in block pellet runs (**Fig. 5M**) on the rPR task resulting in a higher number of pokes per pellet (**Fig. 5N**), suggesting that SK3 over-expression normalizes SNI-induced changes in reward-seeking. Interestingly, these mice earned fewer pellets (**Fig. 5L**), which was due to continuous engagement with the task without resets, leading to longer times between pellets at high poke to reward ratios. Locomotor activity during the rPR task (**Fig. 5I**) and on an open field task (**Fig. S5F**) were not different between groups, arguing against confounding motor effects contributint to behavioral differences. Additionally, we found no differences in a classical anxiety measure, time spent in the center of an open field chamber (**Fig. S5D-E**).

**Figure 5.**
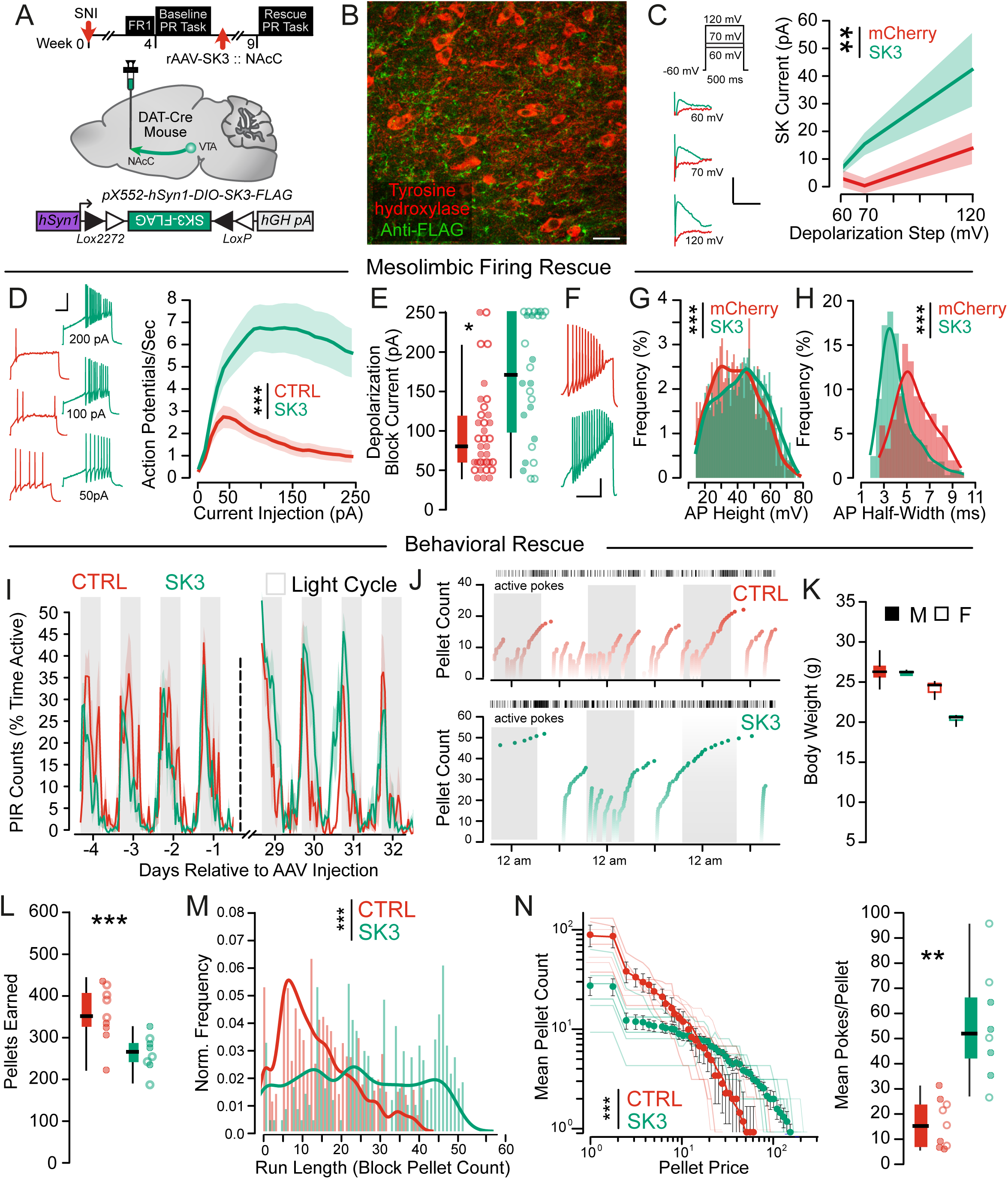
Increasing SK-channel expression rescues firing properties of mesolimbic dopamine neurons and reward seeking following chronic SNI. (A) Timecourse and experimental schematic showing Cre-dependent viral expression of SK3 in mesolimbic dopamine neurons. (B) Histological validation of SK3 virus (anti-FLAG tag) with tyrosine hydroxylase in mesolimbic dopamine neurons (scale bar = 15 µm). (C) SNI neurons had increased SK channel currents (F_Group_ = 9.34, p_Group_ = 0.0034) following injection of an SK3-expressing AAV (n_mCherry_ = 5M/6F, N_mCherry_ = 3M/3F, n_SK3_ =4M /7F, N_SK3_ = 2M/2F). Representative traces (inset, scale bars: 100 pA, 10 ms). (D) SNI neurons (n_mCherry_ = 19M/16F, N_mCherry_ = 9M/7F, n_SK3_ = 11M/15F, N_SK3_ = 3M/3F) had increased action potential firing (F_Group_ = 505.31, p_Group_ < 0.001) following SK3 overexpression. Representative traces are shown (Inset; scale bar = 20 mV, 500 ms). (E) Neurons overexpressing SK3 entered into depolarization block at higher injection currents relative to mCherry expressing controls (t = 3.72, p < 0.001). Males: closed circles, females: open circles. (F) Representative traces of ramp current injections; scale bar = 20 mV, 500 ms. (G) Action potential heights were increased (KS = 0.11, p < 0.001) following SK3 overexpression (n_SNI_ = 13M/11F, N_SNI_ = 8M/5F, n_SK3_ = 11M/15F, N_SK3_ = 2M/3F). (H) Action potential half widths were reduced following SK3 overexpression relative to controls (KS = 0.492, p < 0.001). (I) Homecage locomotor activity was not different in SK3 overexpressing mice (F_Preinjection_ = 1.90, p_Preinjection_ = 0.20, F_Postinjection_ = 3.83, p_Postinjection_ = 0.076). (J) Representative examples of SNI mice following injection of a GFP (top) or SK3 (bottom) expressing AAV performing the rPR task over three days, dark cycle is shaded. (K) Body weight is unchanged (F_Timepoint_ = 3.59, p = 0.068) following injection of a GFP (n = 5M/4F) or SK3 (n = 4M/4F) expressing AAV in SNI mice. (L) Mice overexpressing SK3 in mesolimbic DA neurons obtained fewer pellets (t = 2.84, p = 0.01) during the rPR task. Males: closed circles, females: open circles. (M) SK3 overexpressing mice had longer progressive ratio run lengths (KS = 0.38, p < 0.001) relative to controls. (N) SK3 overexpressing mice expended significantly more pokes per pellet earned (F_Group_ = 100.14, p_Group_ < 0.001, t = 3.97, p = 0.0015). Males: closed circles, females: open circles.

## Discussion

The pathogenesis and specific behavioral consequences of dopaminergic dysregulation in chronic pain has been debated for decades. Our study is unique in that we tested multiple behavioral constructs that are relevant to affective symptoms of chronic pain, including reward valuation, flexibility, reward learning, and effort based decision making, which are inferred from performance on tasks that are known to be modulated by DA. Surprisingly, the ability to form action-outcome associations in instrumental tasks, and to update behavior in response to reward history is intact following SNI, arguing that chronic pain is not associated with generalized impairments in DA function. Although this finding challenges the prevailing model of pain-induced hypodopaminergia^14, 34^, it is consistent with our direct assessments of mesolimbic dopamine neuron function. Ours is the first study to selectively record from genetically and anatomically-defined mesolimbic DA neurons, which exhibited comparable spontaneous firing and passive membrane properties in SHAM and SNI mice, indicating that their ability to fire action potentials is intact or even enhanced following chronic pain.

By contrast, SNI mice were slower to initiate new trials in a self-paced foraging task when the reward rate in the environment was high, suggesting an inability to sustain effort across multiple trials. This deficit in sustaining goal-directed behavior across time was even more apparent in the progressive ratio task, where SNI mice adopted a ‘thrifty’ strategy, earning equivalent pellets while disengaging earlier from the task. Our findings build on prior work establishing the role of mesolimbic DA release for signaling the value of work towards a reward^14, 38, 39, 43^, and the necessity of sustained dopamine neuron firing for maintaining progress towards a goal over time and effort delays^14, 39, 43^. Our recordings revealed that mesolimbic DA neurons more readily enter depolarization block and are unable to sustain regular firing, providing a mechanism for these behavioral deficits. We recorded from identified mesolimbic (NAcC-projecting) DA neurons, which are ‘atypical’, with infrequent spontaneous firing and no prominent hyperpolarization-activated (Ih) current^49^. Despite low expression in this ‘atypical’ DA neuron population, SK channels are critical for setting the maximum firing rate^49^, because they are the main contributors to the mAHP, which allows voltage-gated sodium channels to recover from slow inactivation^48-50^. Consistent with this key role, convergent biophysical modeling and experimental recordings established that SK channel currents were reduced following SNI. Virally over-expressing the SK channel selectively in mesolimbic DA neurons restored both the premature entry of neurons into depolarization block and performance of SNI mice in the PR task. While our model also predicted a small decrease in the A-type potassium current, we did not see significant differences in our direct measurements of the A-type current, and CRISPR-mediated editing of the Kv4.3 channel^52^ did not rescue behavioral adaptations, pointing the privileged role of SK channels in controlling sustained firing of mesolimbic DA neurons and subsequent behavioral adaptations in chronic pain states.

Critically, acute pain did not drive an avolitional phenotype, nor did acute analgesia reverse the behavioral adaptations once established in the chronic pain state. The electrophysiological and behavioral adaptations were not present acutely (1 week) after injury, underscoring the importance of persistent, ongoing nociceptive input in development of negative affect. The heterogeneous definition of ‘chronic’ in preclinical literature (ranging between 2 days and 12 weeks)^53-55^, even within similar pain models, could contribute to variability in results and difficulty in reconciling or translating preclinical findings. Further, mesolimbic DA neurons from SHAM mice exhibited an electrophysiological phenotype that was intermediate between Naive and SNI mice, suggesting that incisional injury may be sufficient to drive persistent adaptations in mesolimbic DA neurons. This graded phenotype argues that the choice of control manipulation may be another important factor contributing to heterogeneity in prior studies seeking to understand mesolimbic DA adaptations in preclinical models.

Two open questions are *why* and *how* these adaptations in mesolimbic DA neurons emerge in chronic pain. While acute pain is adaptive—driving avoidance of injury—persistent alterations in mesolimbic DA signaling may similarly confer an evolutionary advantage by biasing behavior toward energy-efficient strategies that minimize threat exposure in a compromised state. Reduced SK channel function has been observed in amygdala, peripheral and spinal circuits in chronic pain, attributed to downregulated gene expression^56^. Because SK activation is gated by Ca²□ influx through adjacent calcium channels and NMDA receptors^57-59^ and is further tuned by phosphorylation that modulates calcium sensitivity and surface stability^60-63^, multiple mechanisms could plausibly converge to reduce SK function in mesolimbic dopamine neurons. While future work will be needed to elucidate the mechanisms underlying reduced SK channel function, our results nonetheless point to SK channels as a promising therapeutic target for affective symptoms of chronic pain. Importantly, FDA-approved medications including Riluzole and Chlorzoxazone act in part through SK channel modulation^64-66^, providing an immediate translational path toward targeting the affective symptoms of chronic pain.

## Conclusion

Together, our findings establish a causal link between SK channel dysfunction, specific deficits in sustained firing of mesolimbic dopamine neurons, and the emergence of an avolitional state in chronic pain. These adaptations are unique to chronic pain states, and provide a framework and mechanism for reconciling prior paradoxical reports of a pain-related hypodopaminergic state in preclinical models.

## Figure Legends

**Supplemental Figure 1. Acute pain and analgesia do not affect strategy on the progressive ratio task.**

(A) Schematic of acute incisional pain experiment; body weights were not different pre and post paw incision (Pre = 23.04 ± 0.919 sec, Post = 22.56 ± 0.904, F = 0.134, p = 0.717, n=6M/6F).

(B) There was a significant effect of time on homecage activity levels, with slight reductions in overall activity levels following incisional pain (F = 7.01, p < 0.001).

(C) Mice exhibited significantly greater numbers of active FED responses immediately following paw incision (F = 1.57, p = 0.002).

(D-E) There was no difference in pellets earned (F_Group_=1.11, p=0.305, Pre = 378.58 ± 24.11, Post = 345.5 ± 25.99) or poke accuracy (F_Group_=0.01, p=0.948, Pre = 97.3 ± 4.90 sec, Post = 97.4 ± 4.34) before and after paw incision.

(F-G) There was a shift in the distribution of run lengths towards intermediate run lengths (KS: 0.128, p = 0.002), with no differences in demand curves or pokes per pellet before and after acute incisional pain (F_Group_=1.89, p=0.184, Pre = 19.47 ± 4.396, Post = 13.56 ± 2.371).

(H) Experimental design; MET and VEH injections were administered at 6 pm, prior to onset of progressive ratio testing; there was a significant increase in the withdrawal ratio after metformin injection compared to vehicle injection (F = 87.10, p < 0.0001, n= 6M/8F mice).

(I-J) MET had no significant effect on total ambulatory activity (F_Group_ = 0.308, p = 0.579) or on active FED responses (F_Group_ = 0.858, p = 0.357); mean of contemporaneous SHAMs are indicated in grey.

(K) There were no differences in body weight between VEH and MET conditions (F_Group_=0.75, p=0.390, VEH = 24.40 ± 0.62 g, MET = 25.18 ± 0.66).

(L-M) MET did not affect the number of pellets earned (F_Group_=0.25, p=0.620, VEH = 38.14 ± 2.035, MET = 36.64 ± 2.085) or response accuracy (F_Group_=0.37, p=0.549, VEH = 94.19 ± 0.94, MET = 95.26 ± 1.427).

(N-O) Relative to VEH, mice treated with MET did not exhibit a shift in distributions of run lengths (KS: 0.16, p = 0.915) nor a significant difference in pokes per pellet (F_Group_=0.11, p=0.743, VEH = 12.12 ± 2.019, MET = 11.23 ± 1.619).

**Supplemental Figure 2. SNI-associated behavioral adaptations in reversal learning and progressive ratio tasks are emulated by systemic haloperidol.**

(A) Experimental schematic; HAL and VEH injections were administered at 6 pm, prior to onset of progressive ratio testing (n= 10M/10F mice).

(B-C) Low dose HAL did not blunt total ambulatory activity (F_Group_=0.01, p=0.912) but significantly reduced active responses in the progressive ratio task (F_Group_=36.86, p<0.001).

(D) There was no difference in body weight between VEH and HAL conditions (F_Group_=0.03, p=0.862, VEH = 24.18 ± 0.30 g, HAL = 24.29 ± 0.31).

(E-F) HAL did not affect the number of pellets earned (F_Group_=1.24, p=0.273, VEH = 42.44 ± 3.99, HAL = 45.00 ± 2.38) or response accuracy (F_Group_=0.78, p=0.381, VEH = 95.19 ± 1.11, HAL = 93.23 ± 1.84).

(G-H) Relative to VEH, mice treated with HAL exhibited a shift in distributions of run lengths (KS: 0.26, p = 0.028) and a significant reduction in pokes per pellet (F_Group_=9.07, p=0.005, VEH = 21.92 ± 4.61 g, HAL = 7.67 ± 0.89).

(I)There was no difference in pellets earned in the probabilistic reversal task (VEH = 126.00 ± 9.82, HAL = 116.92 ± 13.55, F = 0.32, p = 0.577, n = 6M/7F)

(J-L) There was no difference in Win-stay (VEH = 74.91 ± 2.12, HAL = 75.24 ± 1.99, F = 0.013, p = 0.911) or in Lose-shift (VEH = 45.34 ± 1.63, HAL = 42.91 ± 2.04, F = 0.861, p = 0.363) strategy or on Accuracy (VEH = 58.73 ± 1.99, HAL = 54.68 ± 1.98, F = 2.09, p = 0.161) on the probabilistic reversal task between VEH and HAL-injected mice.

(M) Logistic regression revealed significant decay of weighting of rewarded (F_Trial_=31.47, p<0.001) and unrewarded (F_Trial_= 6.27, p < 0.001) choices over time, but no difference in integration of prior reward history between VEH and HAL-injected mice on the probabilistic reversal task (rewarded: F_Group_ = 0.22, p = 0.64, F_Interaction_= 0.55, p = 0.70, unrewarded: F_Group_ = 1.92, p = 0.17, F_Interaction_= 0.61, p = 0.66).

**Supplemental Figure 3. Paradoxical changes in passive membrane properties of mesolimbic dopamine neurons following SNI.**

(A) Resting potential was unchanged (F_Group_ = 1.37, p_Group_ = 0.26) chronically following SNI (n_NAIVE_ = 4M/17F, N_NAIVE_ = 1M/5F, n_SHAM_ = 16M/24F, N_SHAM_ = 9M/13F, n_SNI_ = 24M/16F, N_SNI_ = 11M/7F). Males: closed circles, females: open circles.

(B) Spontaneous firing was unchanged (H_Group_ = 0.10, p_Group_ = 0.95) at a chronic timepoint after SNI. Males: closed circles, females: open circles.

(C) Input resistance (F_Group_ = 5.85, p_Group_ = 0.004) was increased chronically following SNI (n_NAIVE_ = 6M/18F, N_NAIVE_ = 3M/4F, n_SHAM_ = 16M/24F, N_SHAM_ = 9M/12F, n_SNI_ = 24M/17F, N_SNI_ = 11M/7F). Males: closed circles, females: open circles.

(D) Sag potential mediated by the Ih current during a -100 pA current step was unchanged in SNI neurons (F_Group_ = 0.14, p_Group_ = 0.87) 5 weeks following SNI (n_NAIVE_ = 1M/6F, N_NAIVE_ = 1M/2F, n_SHAM_ = 9M/22F, N_SHAM_ = 4M/10F, n_SNI_ = 22M/5F, N_SNI_ = 11M/4F). Males: closed circles, females: open circles.

(E) Resting potential was unchanged (F_Group_ = 1.20, p_Group_ = 0.31) acutely following SNI (n_NAIVE_ = 4M/17F, N_NAIVE_ = 1M/5F, n_SHAM_ = 16M/7F, N_SHAM_ = 6M/2F, n_SNI_ = 28M/16F, N_SNI_ = 12M/6F). Males: closed circles, females: open circles.

(F) Spontaneous firing was increased in sham neurons (H_Group_ = 8.61, p_Group_ = 0.013) compared to naive and SNI neurons at an acute timepoint after SNI. Males: closed circles, females: open circles.

(G) Input resistance was increased in SNI neurons (F_Group_ = 5.24, p_Group_ = 0.008) 1 week following SNI (n_NAIVE_ = 6M/18F, N_NAIVE_ = 3M/4F, n_SHAM_ = 16M/3F, N_SHAM_ = 6M/1F, n_SNI_ = 23M/5F, N_SNI_ = 10M/2F). Males: closed circles, females: open circles.

(H) The sag potential was unchanged (F_Group_ = 0.0012, p_Group_ = 0.99) by acute SNI (n_NAIVE_ = 1M/6F, N_NAIVE_ = 1M/2F, n_SHAM_ = 7M/3F, N_SHAM_ = 2M/1F, n_SNI_ = 22M/3F, N_SNI_ = 10M/2F). Males: closed circles, females: open circles.

**Supplemental Figure 4. Changes in A-type potassium currents do not account for biophysical adaptations in mesolimbic DA neurons or behavior following SNI.**

(A) SNI neuron models had significantly larger A-type potassium conductance (1.832 ± 0.194 mS/cm^2^) than SHAM (1.565 ± 0.215 mS/cm^2^) neuron models (N_SNI_ = 50, N_SHAM_ = 64, U = 1977, p = 0.032)

(B) The time until the first action potential was increased following SNI (n_SNI_ = 19M/16F, N_SNI_ = 9M/7F, n_SHAM_ = 16M/20F, N_SHAM_ = 9M/11F, F_Group_ = 77.11, P < 0.001)

(C) The A type potassium current was unchanged following SNI (n_SHAM_ = 10M/8F, N_SHAM_ = 6M/5F, n_SNI_ = 7M/9F, N_SNI_ = 3M/4F, t = 0.41, p = 0.61).

Representative traces showing predrug (blue/red), post-AmmTx3 (gold) and the subtracted trace (black) are shown. Scale bars, 50 ms, 1000 pA. Males: closed circles, females: open circles.

(D) Time course and schematic of CRISPR-mediated editing of Kcnd3 (Kv4.3) isoform in mesolimbic dopamine neurons following SNI.

(E) Confocal image of VTA (20x) showing co-localization for SaCas9 and tyrosine hydroxylase in mesolimbic dopamine neurons.

(F) There was a continued decrease in run lengths following genetic editing of Kv4.3 (KS: 0.21, p < 0.001), the predominant subunit contributing to the A-type current in mesolimbic DA neurons.

**Supplemental Figure 5. Exploratory behaviors are unaffected by SK3 overexpression.**

(A) SK3 expressing mesolimbic dopamine neurons had no change in rheobase relative to mCherry-expressing neurons (SNI = 16.9 ± 2.4, SK3 = 16.2 ± 3.3, t = 0.18, p = 0.86). Males: closed circles, females: open circles.

(B) SK3 expressing mesolimbic dopamine neurons had higher peak firing rates relative to mCherry-expressing neurons (SNI = 4.5 ± 0.55, SK3 = 9.85 ± 1.21, t = 4.05, p = 0.0003). Males: closed circles, females: open circles.

(C) SNI neurons had lower (KS = 0.48, p < 0.001) interspike intervals at their peak firing rate following SK3 overexpression.

(D) Representative heatmaps of mouse position during open field test.

(E) There was no difference (T = 0.90, p = 0.38) in time spent in center zone of the arena between control (774.7 ± 72.7, n = 5M/5F) and SK3 (877.3 ± 88.7, n = 4M/4F). Males: closed circles, females: open circles.

(F) Ambulation during an open field task in SK3 overexpressing mice (n_GFP_ = 5M/5F, n_SK3_ = 4M/4F) was unchanged (SK3 = 7430 ± 538, GFP = 7975 ± 1168, t = -0.390, p = 0.7020)

## Methods

### Subjects

Adult male and female mice (6 weeks of age or older) on a C57BL/6J background were used for all experiments. Mice were either obtained directly from Jackson Laboratory, or bred in-house from stock lines obtained from the Jackson Laboratory (DAT-Cre JAX# 027395, or wild type C57BL/6J). Mice were group housed (2–5 per cage, single sex) in a vivarium that was controlled for humidity, temperature and photoperiod (12:12; light onset/offset at 07:00 and 19:00, the light cycle was reversed for photometry recording experiments where indicated). Mice had ad libitum access to food and water outside of FED3.0 experiments. All procedures were approved by the Institutional Animal Care and Use Committee at Washington University in St Louis were conducted in accordance with the Guide for the Care and Use of Laboratory Animals, as adopted by the NIH.

### Drugs

Haloperidol (0.2 mg/kg, i.p.; Sigma-Aldrich, St. Louis, MO, USA) was dissolved in methanol into 1 mg/ml stock solution and diluted to 0.02 mg/ml working solution with 0.9% saline a day before behavioral testing. Metformin (200 mg/kg, i.p.; TCI America, Portland, OR, USA) was dissolved in 0.9% saline a day prior to injection in volumes of 300 μl. Both drugs were administered intraperitoneally. For analgesic screening, von Frey was conducted 30 minutes after metformin injection. For operant behavioral testing, haloperidol and metformin were injected one hour prior to the onset of the dark cycle, and only 7h of data following injection was included in the analyses.

### Stereotaxic surgery

Anesthesia was induced and maintained with isoflurane at 5% and 1.5%, respectively. Mice were placed in a stereotaxic frame and craniotomies were performed. For electrophysiology, photometry and behavioral experiments, purified and concentrated AAVs were injected into the NAcC (coordinates relative to bregma: anteroposterior (AP), +1.6 mm; mediolateral (ML), ±0.9 mm; dorsoventral (DV), −4.0 mm) and the VTA (coordinates relative to Bregma: AP, −3.0 mm; ML, ±0.55 mm; DV, −4.5 mm). Injections were carried out using a Drummond Scientific Nanoject III and graduated pipettes with a tip diameter of 10–15 μm at a rate of 0.05 μl/min for a total volume of 0.25–0.5 μl in each hemisphere; pipettes were left *in situ* for an additional 2 min to allow for the diffusion of virus. The minimum viral expression time was 28 days. For photometry experiments, optic fibers were implanted unilaterally over the right NAcC and secured with dental cement. Animals recovered for at least 1 week before behavioral training, and only mice with verified infection sites and fiber placements were included in the analyses.

### Injury Model Induction

#### Spared Nerve Injury

Anesthesia was induced and maintained with isoflurane at 5% and 2%, respectively. Using sterile technique, the left sciatic nerve was bluntly dissected from surrounding structures at its separation into the tibial, common peroneal, and sural nerves. Taking care to avoid contact with the sural nerve, 6-0 silk thread was placed around the common peroneal and tibial nerves, which were then tightly ligated. A 1mm section of the nerves distal to the ligation was excised and skin was closed with surgical clips. Antibiotic ointment was applied to the incision and the mice were monitored following surgery for signs of distress. Sham animals received the same surgical procedure including exposure of the sciatic nerve, but did not have nerve ligation. Naive animals did not have any surgical manipulation of the hindpaw.

#### Paw Incision Injury

Anesthesia was induced and maintained with isoflurane at 5% and 2%, respectively. Using sterile technique, a 5 mm longitudinal incision of the skin and fascia of the plantar aspect of the hindpaw was made to a depth of 1mm, after which the wound was closed with surgical adhesive.

### Von Frey Assay

Mechanical sensitivity was assessed using an electronic von Fey (eVF) anesthesiometer (Alemo 2390–5; IITC, Woodland Hills, CA) as described previously^67^. Mice were acclimated individually on an elevated wire mesh grid in a 4” x 4” x 5” plexiglass cylinder for one hour. A semi-flexible tip connecting to the eVF was used to apply pressure to the lateral aspect of the plantar (the sural nerve innervating territory) of the hind paw for both legs. The maximum pressure applied before the animal withdrew its paw was recorded. Stimulation on each paw was separated by a minimum three-minute interval. Five trials of paw withdrawal were conducted on both the left and right hind paws. The maximum and minimum thresholds recorded were excluded from the analysis. The remaining three withdrawal thresholds were used to calculate the average thresholds for both sides. A withdrawal ratio was calculated by dividing the average withdrawal threshold of the injured side (right) by that of the uninjured side (left).

### FED-based operant behavior

FED3.0 devices (OpenEphys) were used for behavioral tasks^35^; mice had constant access to the device which dispensed standard grain pellets (Bio-Serv: Dustless Precision Pellets, Flemington, NJ, USA) as the only source of nutrition available. Mice underwent at least 2 nights of initial FR1 operant training; one nosepoke was ‘active’ and pokes resulted in a 200 ms auditory tone followed by a pellet dispense; the ‘inactive’ poke had no consequence. Body weights were monitored daily at 5 pm for all behavioral assays, any mice who dropped to 85% of their body weight were excluded from further testing.

#### Deterministic Reversal Learning (100:0)

All mice began with the left nosepoke as the ‘active’ poke; an active poke resulted in 1 pellet being dispensed; additional pokes on the active port had no consequence until the pellet was removed from the well. ‘Inactive’ pokes resulted in a distinct error tone and a 2 second time out, during which additional pokes reset the 2 second time out period. Once mice earned 30 pellets, the active and inactive ports switched. The first 72 hours of task performance was included in the analysis.

#### Probabilistic Reversal Learning (80:20)

All mice began with the left nosepoke as the ‘High probability’ poke, which lead to a pellet dispense on 80% of trials (chosen by random number generator in the FED), and lead to a 2 second timeout on 20% of trials. Pokes on the “Low probability’ poke lead to a pellet dispense on 20% of trials, and to a 2 second timeout on 80% of trials. Once mice earned 30 pellets, the active and inactive ports switched. The first 72 hours of task performance was included in the analysis.

#### Dynamic Foraging Task

All mice began with the left nosepoke as the “High probability’ poke, the right as the ‘Low probability’ poke. Every 30 pellets the ‘High’ and ‘Low’ probability assignments switched; however, the probabilities at the beginning of each block were drawn randomly to be a ‘High Reward Rate - 90:60’ (High Probability Poke = 90% pellet, 10% time out, Low Probability Poke = 60% pellet, 40% time out), ‘Medium Reward Rate - 65:35’ (High Probability Poke = 65% pellet, 35% time out, Low Probability Poke = 35% pellet, 65% time out) or ‘Low Reward Rate - 40:10’ (High Probability Poke = 40% pellet, 60% time out, Low Probability Poke = 10% pellet, 90% time out). The first 240 hours of task performance is included in the analysis. Trial initiation latency within a bout (< 15 seconds between successive pellets) was calculated as the interval between a successful trial and the subsequent trial, minus the pellet retrieval time.

Logistic regression was applied to predict each mouse’s choice as a function of outcome history. The model uses the rewarded choice history *R*(*i* - *j*) (equation 1) and the unrewarded choice history *N*(*i* - *j*) (equation 2) to predict choice 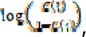 where *C*(*t*) = 1 for a left choice and 0 for a right choice. *R* = 1 for a rewarded choice and 0 for an unrewarded choice, and *N* = 1 for an unrewarded choice and 0 for a rewarded choice. A positive regression coefficient suggests the tendency to repeat previous action, while a negative regression coefficient indicates the likelihood to switch to the other lever. Logistic regression was fit using maximum likelihood estimation in Python’s Statsmodels.

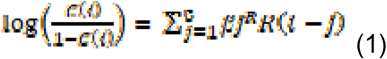

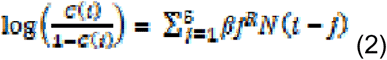

#### Progressive Ratio Task

One nosepoke was designated “Active”; with each successive pellet, the required number of active pokes increased according to the schedule:

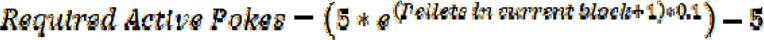

If no pokes were logged within a 30 minute period, the required number of active pokes reset to 1. Mice performed the task for 7 days and were checked each day to ensure adequate nutrition and device function. The final 72 hours of the task were used for analysis to ensure task acquisition. Custom Python scripts were used to extract the total active pokes, total number of pellets delivered, accuracy (active pokes divided by total pokes), and pokes per pellet (total left pokes divided by total pellets), Block pellet count (maximum number of pellets delivered in a run until the 30 minute inactivity reset) and demand curves for group comparisons during the task.

### Patch clamp electrophysiology

Horizontal brain slices (210 μm) were prepared in ice cold choline chloride cutting solution containing the following (in mm): 110 choline chloride, 25 glucose, 25 NaHCO3, 7 MgCl2, 11.6 ascorbic acid, 3.1 sodium pyruvate, 2.5 KCl, 1.25 NaH2PO4, and 0.5 CaCl2, bubbled with 95% O2 and 5% CO2. Slices were kept at 28–30°C in a recording chamber superfused with 2.5 ml/min artificial CSF. For all intrinsic excitability metrics, the external recording solution contained (in mm): 119 NaCl, 2.5 KCl, 1.3 MgCl, 2.5 CaCl2, 1.0 Na2HPO4, 26.2 NaHCO3, and 11 glucose. Fluorescent cells were targeted for recording using an Olympus B50X upright microscope visualized with either 470 nm or 550 nM light for EGFP or mCherry, respectively. Pipette resistance was between 2.8 and 4 MΩ and contained (in mm): 130 potassium gluconate, 10 creatine phosphate, 4 MgCl2, 3.4 Na2ATP, 0.1 Na3GTP, 1.1 EGTA, and 5 HEPES + 5 mg/mL Biocytin. Slices were recovered and immunostained for tyrosine hydroxylase and calbindin in a subset of experiments. For all recordings, resting potential was maintained at −60 mV; recordings were discarded if the access resistance varied by >20%.

Input-Output curves were generated by measuring the action potentials generated in response to 2000 ms current injections over a range of 0-250 pA in 10 pA intervals. Rheobase, peak firing frequency, depolarization block current, spike latency and mAHP were measured from this protocol. Depolarization block current was defined as a decrease in firing rate of at least 25% after the peak that was sustained for at least 3 sweeps. AP half-width was calculated from the current step 2x greater than the current step which elicited the maximum number of action potentials. Half-widths greater than 10 ms were excluded as these likely represent non-physiologic signals. Input resistance was calculated by averaging the membrane voltage change in response to 20 successive injections of -5 pA. Phase plots and action potential threshold were based on the average of the first action potential generated in response to each of 5 successive identical ramp current injections (500 pA over 1000 ms). Data was analyzed using custom Python scripts.

#### A-type measurements

The A-type potassium current was evoked in voltage clamp from a holding potential of -60 mV using voltage steps to -108 for 25 ms, -28 mV for 100 ms, -48 mV for 200 ms, and back to the holding potential every 5 s for 5 trials^52^. A-type potassium current was calculated as the peak minus the steady state of the trace formed after subtraction after the bath application of AmmTx3 (10 nM, Alomone Labs).

#### SK-measurements

The SK current was evoked in voltage clamp from a resting potential of - 60 mV using 100 ms duration depolarizing voltage steps to 0 mV, 10 mV, and 60 mV separated by 4 s for 3 trials. The SK-mediated component of the I_AHP_ was defined as the peak of the outward current following subtraction of the recording after the bath application of apamin (100 nm, Alomone Labs). For experiments using synaptic stimulation, SCH 39166 (1 µm, Tocris), Sulpiride (1 µm, Tocris), LY 341495 (1 µm, Tocris), Picrotoxin (1 µm, Tocris), and CGP 55845 (1 µm, Tocris) were added to the aCSF for blockade of D1, D2, group 2 metabotropic glutamate, GABA_A_, and GABA_B_ receptors, respectively^68^. Experiments were done in current clamp configuration and currents were elicited using a bipolar stimulating electrode placed anterior to the VTA.

### Biophysical modeling

Multicompartment Mesolimbic (VTA→NAc*^DA^) neuron model.* We developed a multicompartment cable model of a mesolimbic (VTA→NAc^DA^) neuron using the NEURON simulation environment (v8.2)^69^ in the Python programming language (v3.10.9)^70^. We implemented a simplified ball-and-stick morphology that affords detailed biophysical simulation while maintaining computational simplicity for high-throughput simulations^71^. We used a soma with a length and diameter of 10 μm, and an axon hillock with a length of 30 μm and a diameter of 0.5 μm. We modeled the dendrite as a single branch with a diameter of 0.5 μm and a length of 1000 μm to reproduce the experimentally measured input resistances of mesolimbic DA neurons. We modeled a myelinated axon with 100 nodes of Ranvier and 99 internode regions^71^.

We included explicit representations of the voltage- and calcium-gated ion channels found in mesolimbic DA neurons as included in the state-of-the-art model of an atypical mesolimbic neuron described and parametrized by Knowlton and colleagues^49^. Specifically, we included explicit representations of the voltage-gated sodium channel Nav1.2, a delayed-rectifier potassium channel, an A-type potassium channel, a small-conductance calcium-activated potassium channel (SK), an N-type voltage-gated calcium channel, an L-type voltage-gated calcium channel, and a linear leak conductance. We also included an explicit representation of a large-conductance calcium- and voltage-gated potassium channel (BK) to better fit our experimentally measured afterhyperpolarization potentials^72^. We expressed these ion channels only in the hillock to reduce dimensionality and therefore computational demand during model population parametrization (see below). We modeled the soma and dendrite as passive cables with only a membrane capacitance and linear leak conductance. The nodes of Ranvier in the axon model contained NEURON’s built-in model of Hodgkin-Huxley nodal dynamics (i.e., the ‘hh’ mechanism in NEURON) to facilitate action potential propagation along the axon. We modeled the internode regions as passive cables (i.e., containing only membrane capacitance and linear leak conductance)^71^.

Parametrizing the mesolimbic (VTA→NAc*^DA^) neuron model populations*. We generated a population of biophysically distinct mesolimbic DA neuron models with variable ion channel expression profiles using Goodman and weare’s Affine-Invariant Markov Chain Monte Carlo method (MCMC)^73^ using the *emcee* Python package as described previously^74^. We used the MCMC method to generate *de novo* combinations of maximal ion channel conductances to model biophysically distinct populations of mesolimbic neuron models. MCMC methods use Bayes’ theorem to estimate posterior probabilities of a given combination of parameter values describing a system based on experimental data and user-defined priors. By separately applying the MCMC method to our patch-clamp data measured from the SNI and SHAM experimental populations, we generated two sets of maximal ion channel conductances that produce mesolimbic neuron models that described SNI and SHAM neuron model populations.

We assumed uniform distributions for each parameter as priors and constrained each distribution to physiologic ranges (e.g., ionic conductances must be greater than or equal to zero). For each class of mesolimbic DA neuron (i.e., SNI and SHAM), we ran the MCMC method ten times, with each successive run injecting progressively increasing variance of Gaussian noise in the initial guess for each parameter’s value to ensure the algorithm fully explored each parameter space. Each run of the MCMC algorithm used 400 independent Markov chains (i.e., ‘walkers’ in the *emcee* package) which iterated their unique combinations of parameters 25 times per run.

Our implementation of the MCMC method functionally serves to systematically generate and test hundreds of thousands of possible combinations of maximal ion channel conductances. Each generated combination of maximal conductances is then used to simulate model behavior with that set of conductances in response to the same current injection protocols used to calculate excitability metrics in our experimental populations (e.g., I/O curve, mAHP, input resistance). We then calculated the normalized Euclidean distance between a given model’s values for each excitability metric and the SNI or SHAM experimental populations’ average values to determine a ‘score’ for that parameter set, where a lower score represents a given parameter set’s validation metrics were closer to the experimentally measured means.

#### Simulation details

For all NEURON simulations, we used a temperature of 30 °C, an initial resting membrane potential of -60 mV, and an integration time step of 25 μs. We calculated each compartment’s time-varying membrane voltage using a backward Euler implicit integration method. For all simulations, we allowed 500 ms prior to current clamp stimulation to ensure each model’s membrane potential reached its equilibrium resting potential prior to perturbation.

### Viral vector and virus generation

The nucleotide sequence of mouse SK3 channel (KCNN3, GenBank AF3577241.1) with a GSSG linker and C-terminal FLAG tag was custom synthesized from Integrated DNA Technologies (Coralville, IA) with 5’ EcoRV and 3’ SpeI sticky ends. We cloned this fragment into pX552:hSyn:DIO:Synaptophysin-GFP^75^ at EcoRV and SpeI sites on either side of the DIO cassette to generate the pX552:hSyn:DIO:SK3-FLAG viral vector. We verified the construct using whole plasmid long-read sequencing (Plasmidsaurus). AAVDJ-serotyped virus (titer 1.64 x 10^13^ vg/ml) was produced by the Hope Center Viral Vector Core. All other viruses: GRAB-DA (pAAV-hSyn-GRAB-gDA3m, #208698-AAV1), retrograde GFP (pAAV-hSyn-DIO-EGFP, #50457-AAVrg), retrograde mCherry (pAAV-Ef1a-fDIO-mCherry, #114471-AAVrg) were obtained from Addgene.

### Immunohistochemistry

For post-hoc analysis of biocytin filled cells following electrophysiological recordings, slices were collected and transferred into 4% paraformaldehyde overnight for fixation. Slices were then moved to PBS and stained within 7 days of fixation. Slices had 3 PBS washes for 10 min each followed by blocking in 10% normal donkey serum with 0.3% Triton-X-100 in PBS for 1 h at room temperature. Slices were then incubated with rabbit anti-calbindin (1:200, Cell Signaling Technology, 13176) and mouse anti-TH (1:500, Sigma-Aldrich, T2928) with 3% normal donkey serum and 0.3% Triton-X-100 for 48 h at 4°C. Slices had 3 washes in PBS for 15 min and were then incubated in goat anti-mouse AlexaFluor 555 (1:1000, Thermo Fisher) and goat anti-rabbit AlexaFluor 488 (1:1000, Thermo Fisher) overnight at 4°C. Slices were mounted, coverslipped, and imaged using a Leica SP5 confocal microscope at 20x.

Brains from mice that underwent behavioral testing with SK3 manipulation were fixed with 4% PFA, sliced at 40 µm and stored in PBS. To verify viral expression, FLAG sequence encoded in the SK3 virus was labeled. Slices had 3 PBS washes for 10 min each followed by blocking in 10% normal donkey serum with 0.5% Triton-X-100 in PBS for 1 h at room temperature. Slices were then incubated with mouse anti-FLAG (1:1000, Millipore F1804, lot#: 0000448665) and rabbit anti-TH (1:1000, Millipore AB152, lot#: 4266791) with 5% normal donkey serum and 0.5% Triton-X-100 for 24 h at 4°C. Slices had 3 washes in PBS with 0.5% Triton for 10 min and were then incubated in donkey anti-mouse AlexaFluor 488 (1:1000, Jackson ImmunoResearch Laboratories 715-545-150, Lot#: 175021) and goat anti-rabbit AlexaFluor 647(1:1000, Invitrogen A21245, lot# 2833435) for 2 h at room temperature. Slices were mounted, coverslipped, and imaged using a Leica DMI8 confocal microscope at 20x.

### Locomotor Activity

Homecages were equipped with a wireless passive infrared (PIR) activity monitor to measure mouse activity levels (Models 4430 and 4610 from MCCI Corporation, Ithaca NY). PIR activity data, and environmental measurements (temp, humidity, light levels) were transmitted via Low frequency Radio Wide Area Network (LoRaWAN) using Internet of Things (IoT) infrastructure (The Things Network, Netherlands), saved in a cloud-database (InfluxDB), and visualized with an online dashboard (Grafana). In a subset of mice, open field activity was collected to assess changes in locomotion and anxiety-like behavior. For 1 hour, mice were placed in a 47.6 x 25.4 x 20.6 cm chamber equipped with a grid of bisecting photobeams (SmartFrame, Kinder Scientific) to automatically track position and behavior.

### Statistical Analysis

Statistical significance was accepted at the level of p < 0.05. Averaged values are presented as means ± SEM. Boxplots show median and interquartile range. Custom Python scripts were used for analysis. For parametric data, one-way or two-way ANOVA with Tukey posthoc tests and unpaired or paired t-tests were used where appropriate. For non-parametric data, the Kruskall-Wallis test with Dunn’s pairwise posthoc testing with the Bonferroni correction was used. For comparisons of distributions, the Kolmogorov-Smirnov (K-S) test was used. A cluster-based permutation test was used to compare action potential phase plots.

## Acknowledgements

We would like to thank members of the Creed lab at Washington University for helpful discussions. We would also like to thank Ming Jie Li and the Hope Center Viral Vectors Core at Washington University School of Medicine, Washington University’s Division of Comparative Medicine Animal Facilities, along with Judith Golden and Catherine Pena for technical help and discussions. This work was supported by the National Institute on Drug Abuse (R01DA049924, R01DA058755, R01DA056829, R01DA0625509 to M.C.C, T32DA007261), National Institute on Neurological Disorders and Stroke (RM1NS135283 to M.C.C. and R01NS130046 to B.A.C, and L70NS134089 to R.D.G) and McDonnell Center for Systems Neuroscience (Pilot grant to R.D.G). JMT was funded by an Allison Cole Endowed Mentored Research Training Grant from the Foundation for Anesthesia Education and Research (FAER) and a physician-scientist T32 award from NIGMS (T32GM108539).

## Author Contributions

J.M.T, J.P.G, Y.H.C, C.X., E.L. performed behavioral experiments, J.M.T, J.P.G. Y.H.C performed surgical manipulations. J.M.T., H.W. and M.C.C. performed electrophysiology experiments. R.D.G. contributed computational modelling. J.M.T, C.X., A.L., R.G., and M.C.C performed data analysis. A.V.K contributed FED3.0 resources, training and task design. V.K. and B.C. designed and validated custom viral preparations. M.C.C., J.M.T., A.V.K and B.C. secured funding to support the work. J.M.T, J.P.G., R.D.G and M.C.C. wrote the manuscript with input from all authors.

