## Supplemental Figures for "Reduced SK channel control of mesolimbic dopamine neuron firing drives reward seeking adaptations in chronic pain"

Supplemental Figure 1. Acute pain and acute analgesia do not induce behavioral adaptations in progressive ratio responding.

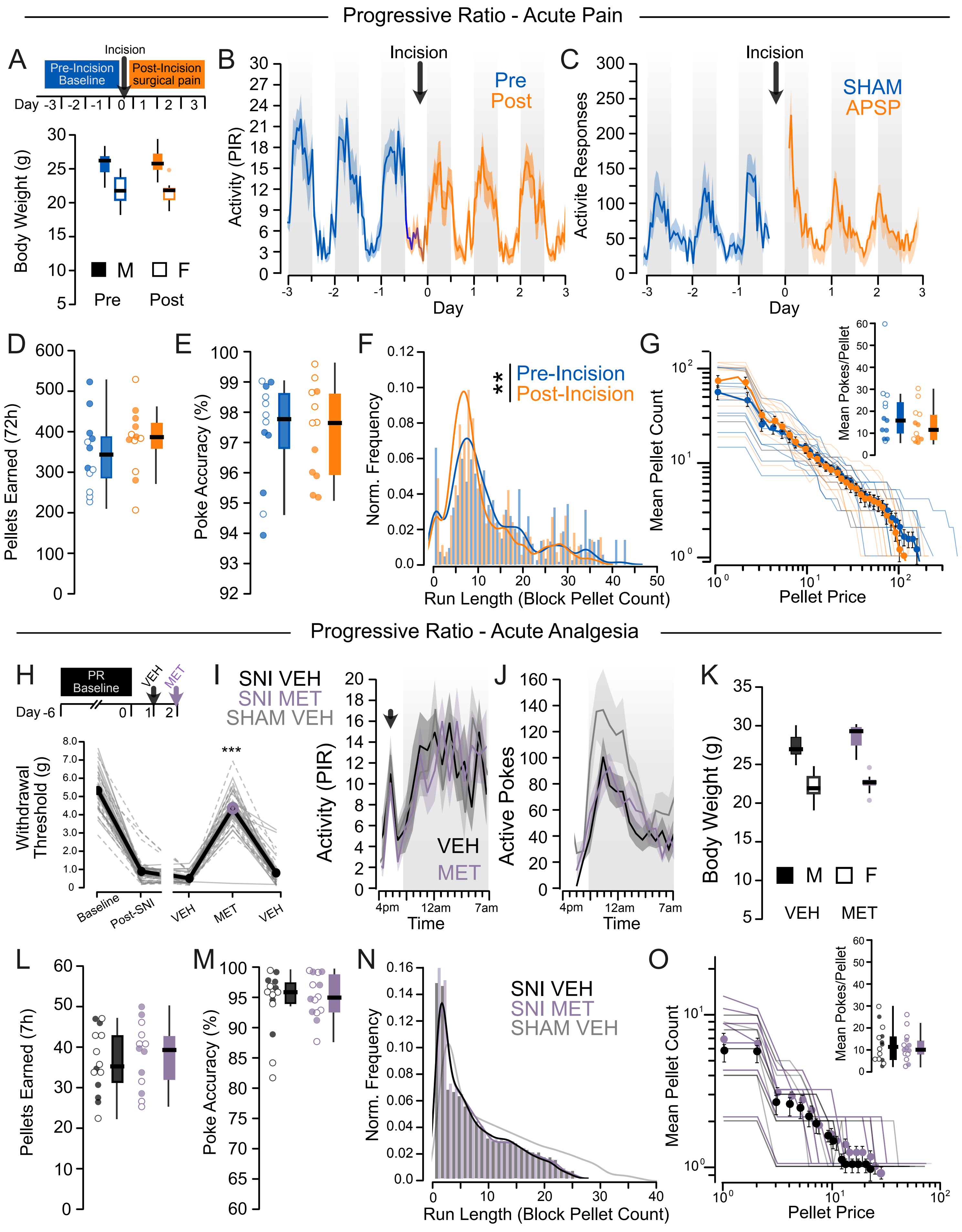

### Supplemental Figure 2 SNI-associated behavioral adaptations in reversal learning and progressive ratio task are emulated by systemic haloperidol.

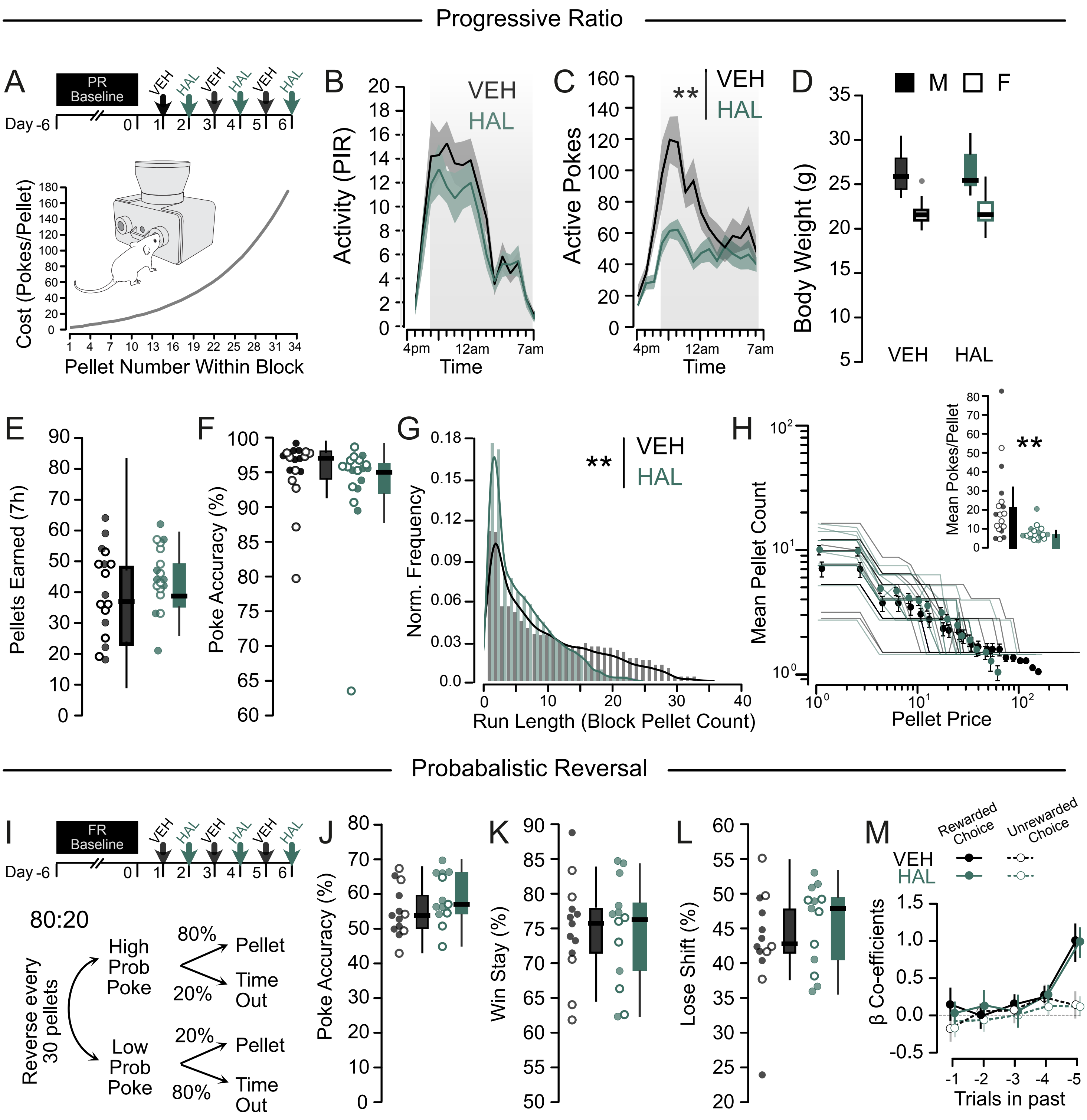

Supplemental Figure 3. Intrinsic membrane properties following SNI.

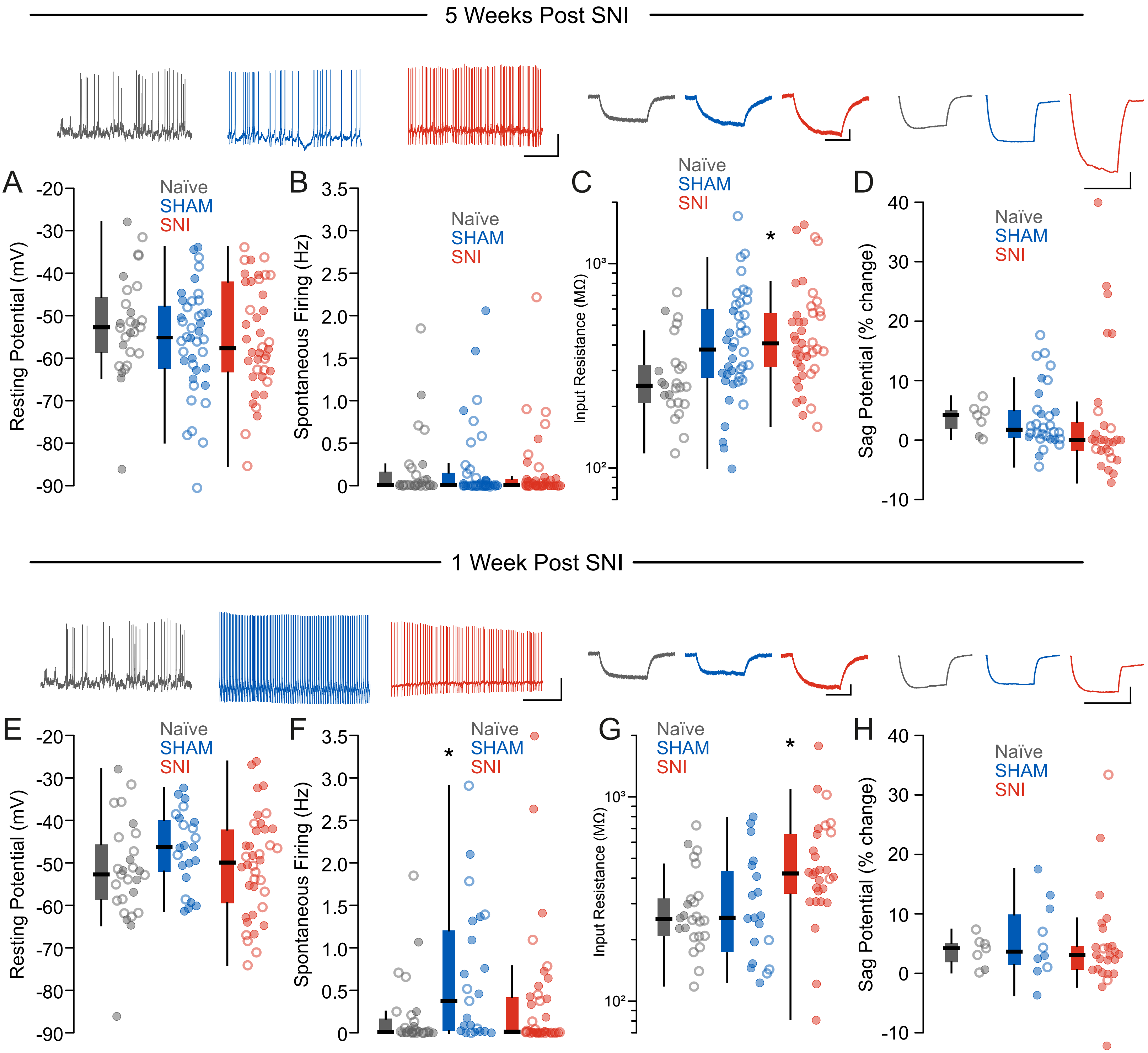

Supplemental Figure 4. Changes in A-type potassium currents do not account for biophysical changes in mesolimbic DA neurons or behavior following SNI.

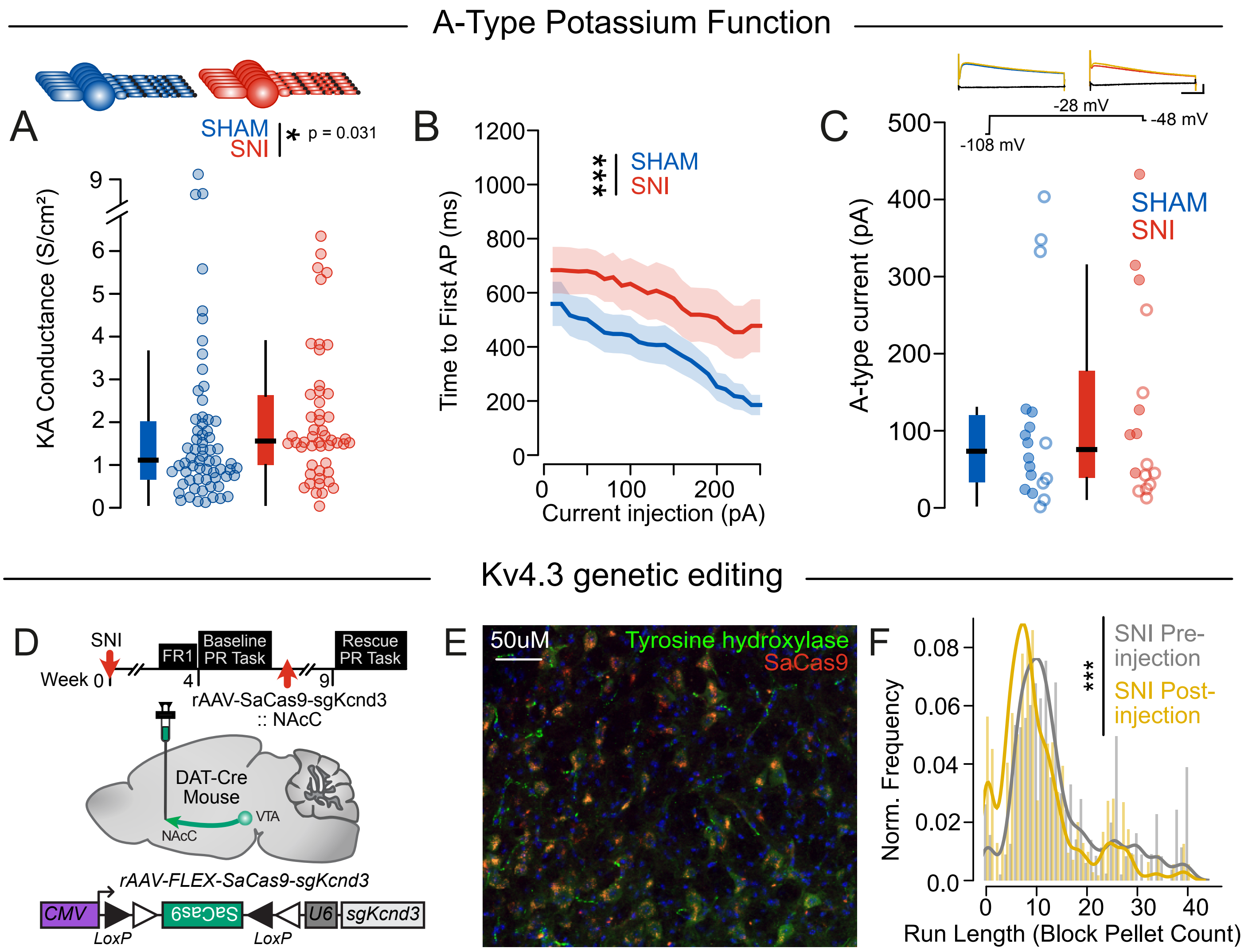

Supplemental Figure 5. Validation of SK overexpression on mesolimbic dopamine neurons and locomotor behavior.

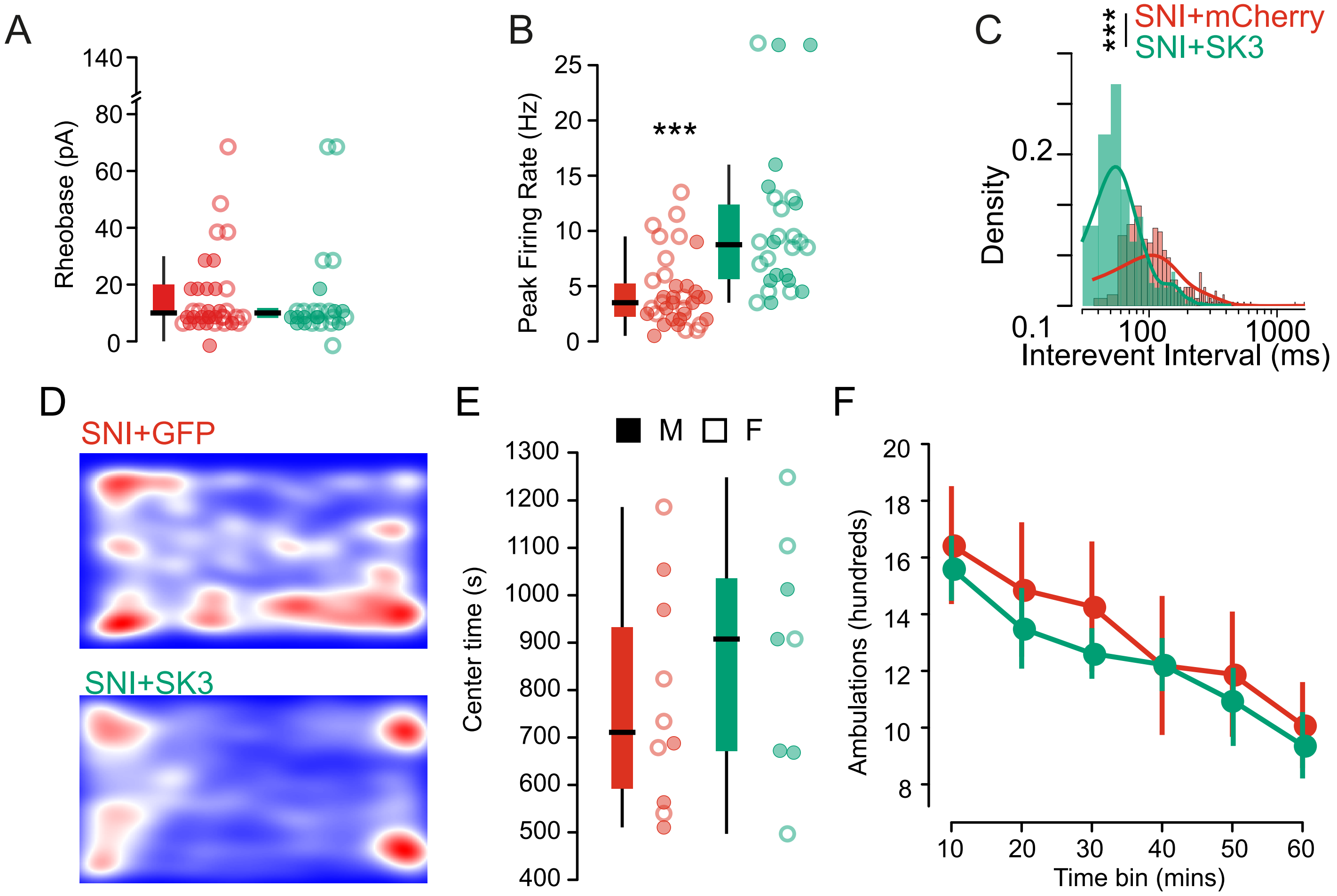
